# Material Properties Of The Embryonic Small Intestine During Buckling Morphogenesis

**DOI:** 10.1101/2024.08.07.606927

**Authors:** Jenny Gao, Lucia Martin, Elise A. Loffet, John F. Durel, Panagiotis Oikonomou, Nandan L. Nerurkar

## Abstract

During embryonic development, tissues undergo dramatic deformations as functional morphologies are stereotypically sculpted from simple rudiments. Formation of healthy, functional organs therefore requires tight control over the material properties of embryonic tissues during development, yet the biological basis of embryonic tissue mechanics is poorly understood. The present study investigates the mechanics of the embryonic small intestine, a tissue that is compactly organized in the body cavity by a mechanical instability during development, wherein differential elongation rates between the intestinal tube and its attached mesentery create compressive forces that buckle the tube into loops with wavelength and curvature that are tightly conserved for a given species. Focusing on the intestinal tube, we combined micromechanical testing with histologic analyses and enzymatic degradation experiments to conclude that elastic fibers closely associated with intestinal smooth muscle layers are responsible for the bending stiffness of the tube, and for establishing its pronounced mechanical anisotropy. These findings provide insights into the developmental role of elastic fibers in controlling tissue stiffness, and raise new questions on the physiologic function of elastic fibers in the intestine during adulthood.

**GRAPHICAL ABSTRACT:** 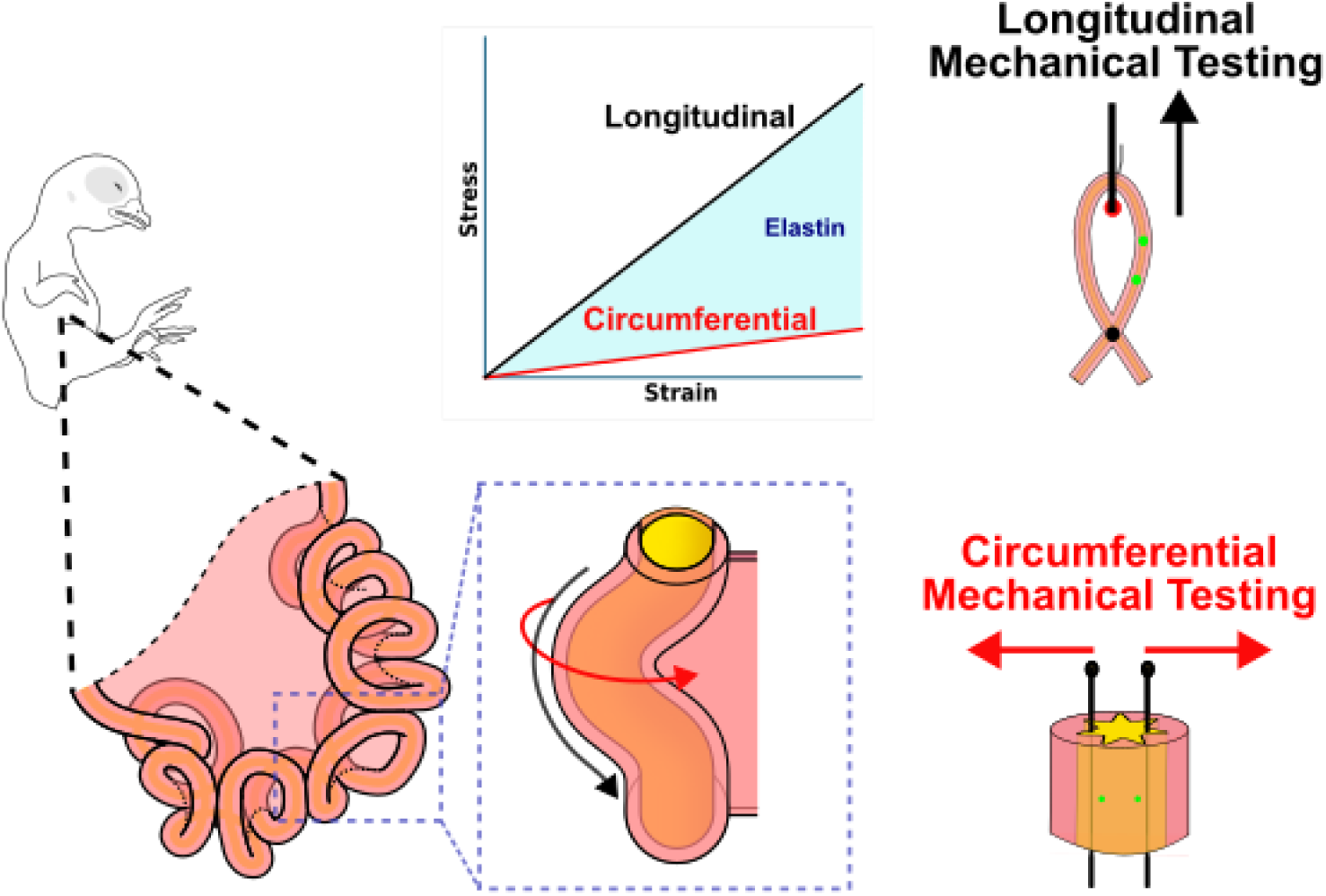

## 1. Introduction

During embryonic development, organs are sculpted from relatively simple origins through precise physical transformations that require stereotyped forces that act on tissues with tightly controlled material properties. In contrast to adult tissues, where the biomechanics field has made great advances in understanding how structure and composition give rise to mechanical function [1], we still understand relatively little about the underpinnings of tissue material properties during embryonic development. As the role of extracellular matrix (ECM) in regulating morphogenesis is increasingly appreciated [2–4], however, there is a growing interest in understanding how physical properties of developing tissues are established [5–7], and how those properties dictate morphological outcomes during organogenesis [8–12]. In the present study, we aim to understand mechanics of the embryonic small intestine, and to relate these properties to the mechanical process by which the lengthy intestinal tube is packed within the body cavity. Errors in this process result in devastating congenital disorders such as midgut volvulus, in which the intestine becomes entangled with itself, obstructing the bowel and strangulating the vasculature [13]. This can result in newborns vomiting bile at birth due to an inability to pass contents through their intestine.

The absorptive function of the small intestine is enabled by its expansive surface area. In addition to villi lining the lumen of the intestine, surface area is maximized by elongation of the intestinal tube, which can extend over 20 feet for humans. Packing of this lengthy tube within the confines of the body cavity without self-entanglement is achieved embryonically through the organization of the tube into undulating loops, a process known as gut looping. Remarkably, the number and curvature of loops is tightly conserved for a given species [8], suggesting that the physical process that transforms the initially straight intestinal tube into its looped form is under regulatory control at the biological level. Looping occurs due to a mechanical instability, generated by the constrained elongation of the intestinal tube against its attached mesentery (Fig. 1). The mesentery is a thin membranous tissue that anchors the intestinal tube to the dorsal body wall. As the embryo develops, both tissues grow, but the intestinal tube elongates much more rapidly than the mesentery, generating tension in the mesentery and compression on the tube. Ultimately, this growth differential becomes so dramatic that compression drives buckling of the intestine into loops [8]. The wavelength and curvature of loops can be faithfully predicted from geometry, growth rates, and stiffnesses of the tube and mesentery, suggesting that each of these properties must be tightly regulated during development to achieve proper looping [12,14].

**Figure 1:**
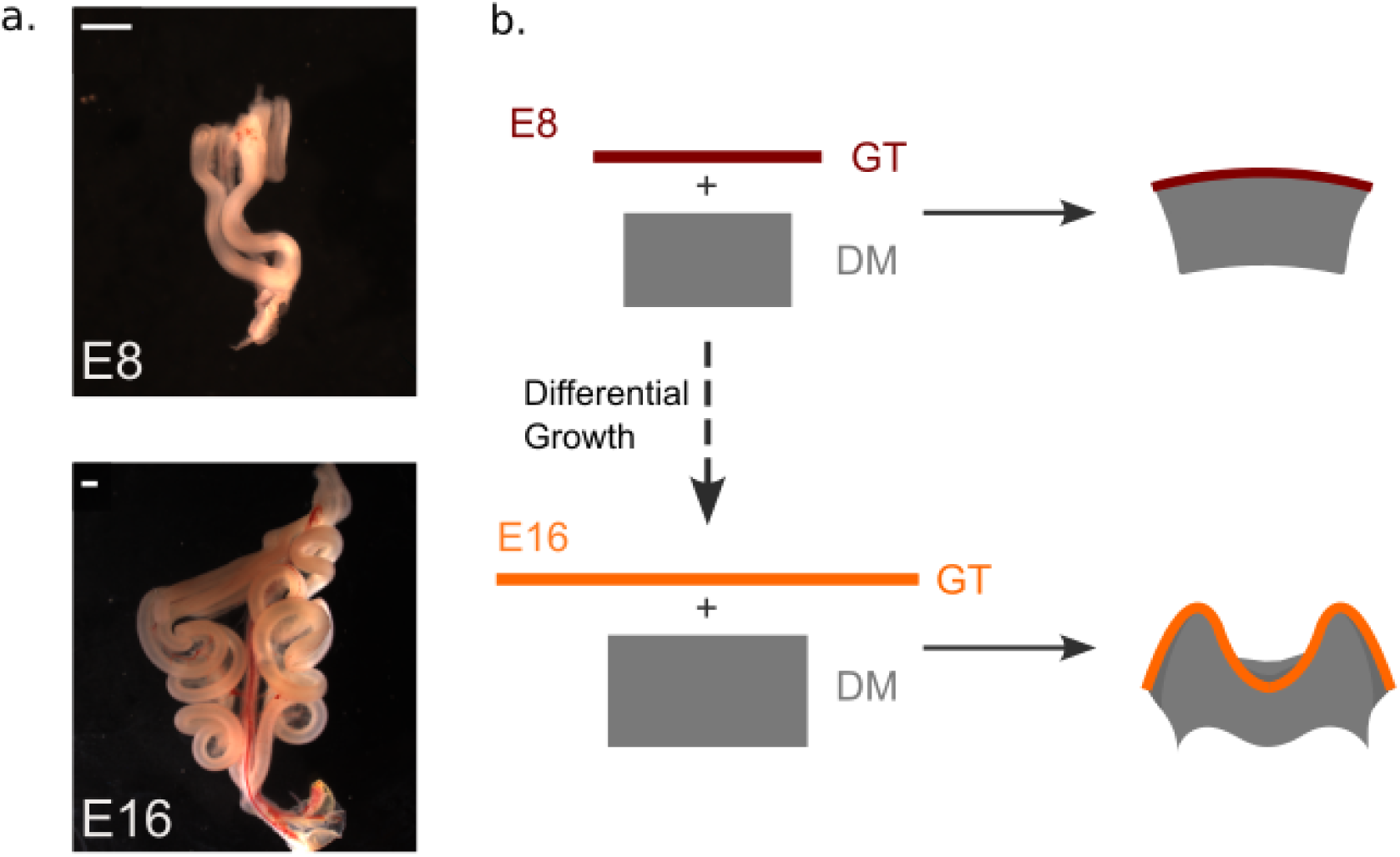
Looping morphogenesis of avian small intestine. (a) The developing small intestine of an embryonic day 8 (E8) and E16 chicken embryo, illustrating the elaboration of many loops as the intestinal tube elongates against its constrained mesentery; scale bar = 1 mm. (b) Schematic description of gut looping by differential growth: the gut tube (GT) and dorsal mesentery (DM, gray) are bonded to one another, and over time, the tube elongates faster than the attached mesentery, creating a growth mismatch that produces stretch in the mesentery and compression on the tube. From the onset of looping at E8 to its completion at E16, the growth differential is exacerbated, resulting in a mechanical instability that forces the intestinal tube into loops.

Increasingly, studies on the mechanics of embryonic development have begun to integrate biophysical and cell biological influences to understand how governing molecular cues specify the forces that sculpt the embryo [9,15–18], and how those forces in turn regulate cell behaviors during development [19–21]. For example, differential growth rates between the rapidly elongating intestinal tube and its constraining mesentery are under control of the Bone morphogenesis protein (BMP) signaling pathway, such that BMP activity limits growth of the mesentery to accentuate differential growth, increasing compressive forces that drive buckling [22]. In order for differential growth to generate sufficient forces to drive buckling, the mesentery and tube must each possess precise material properties as well as growth rates. In recent work, we have identified elastin as a key contributor to mesentery stiffness, with enzymatic depletion of elastin reducing the tensile modulus of the tissue by as much as 90% [7]. Yet, when elastin was removed from the intact intestine at the conclusion of looping by embryonic day (E) 16, no change in loop morphology was observed. This could be explained if elastin depletion similarly reduces both mesentery and tube stiffness, as the buckled morphology depends on the ratio of modulus between these tissues [8]. However, the determinants of material properties in the developing intestinal tube - like most embryonic tissues - are not well understood. In the present study, we investigated the stage-dependent tensile mechanics of the embryonic chick small intestine, revealing pronounced anisotropy that is attributed almost entirely to the elastic ECM of the tube. Understanding embryogenesis from the standpoint of material properties provides new insights into morphogenesis as well as the developmental origins of the vital mechanical functions carried out by tissues and organs during adulthood. In turn, this will advance our understanding of congenital and adult onset diseases, respectively.

## 2. Materials and Methods

### 2.1 Embryos

Fertilized White Leghorn chicken eggs (University of Connecticut Poultry Farm) were incubated at 37° C and 60% humidity in a forced air incubator. E8, 10, 12, 14, and 16 embryos were sacrificed by decapitation, and dissected in PBS to isolate the intestinal tube for histology and mechanical testing.

### 2.2 Longitudinal testing of intestinal tube mechanics

Longitudinal mechanical properties of the intestinal tube were measured in uniaxial tension using a custom micromechanical tester that sits atop the stage of a Zeiss Axiozoom fluorescent macroscope, as described in Loffet et al. [7]. Briefly, 2 cm long sections of intestinal tube were dissected from the intact intestine after removal of the mesentery. The ends of the samples were then secured into PDMS-coated plates and hooked to a fine tungsten cantilever that was displaced by a linear actuator(Fig. 2a) allowing it to apply stretch to the tissue sample. The cantilever was displaced at a rate of 0.5% of tissue length per second. Cantilever bending stiffness (0.45 mN/mm for E12 and E16 samples; 0.07 mN/mm for E8 samples) was used to compute applied force from beam deflection. Cross sectional area was determined from cryosections of stage-matched intestinal tissue. Particles of lipophilic fluorescent vital dye DiO were placed on the tissue to serve as fiducial markers for strain calculation; the cantilever tip was also identified by fluorescent marking. Automated tracking of tissue displacements and cantilever deflection were conducted on time lapse movies of mechanical tests by particle tracking using custom MATLAB code, and used for subsequent calculation of strain and force, respectively. For simplicity, tissue deformations were quantified as linear/engineering strain as the change in distance between DiO labels divided by the undeformed distance between them. Stress was quantified as the force divided by the undeformed cross-sectional area. Finally, elastic modulus was quantified by least squares curve fitting of the stress-strain curve, which was linear for all samples analyzed.

**Figure 2:**
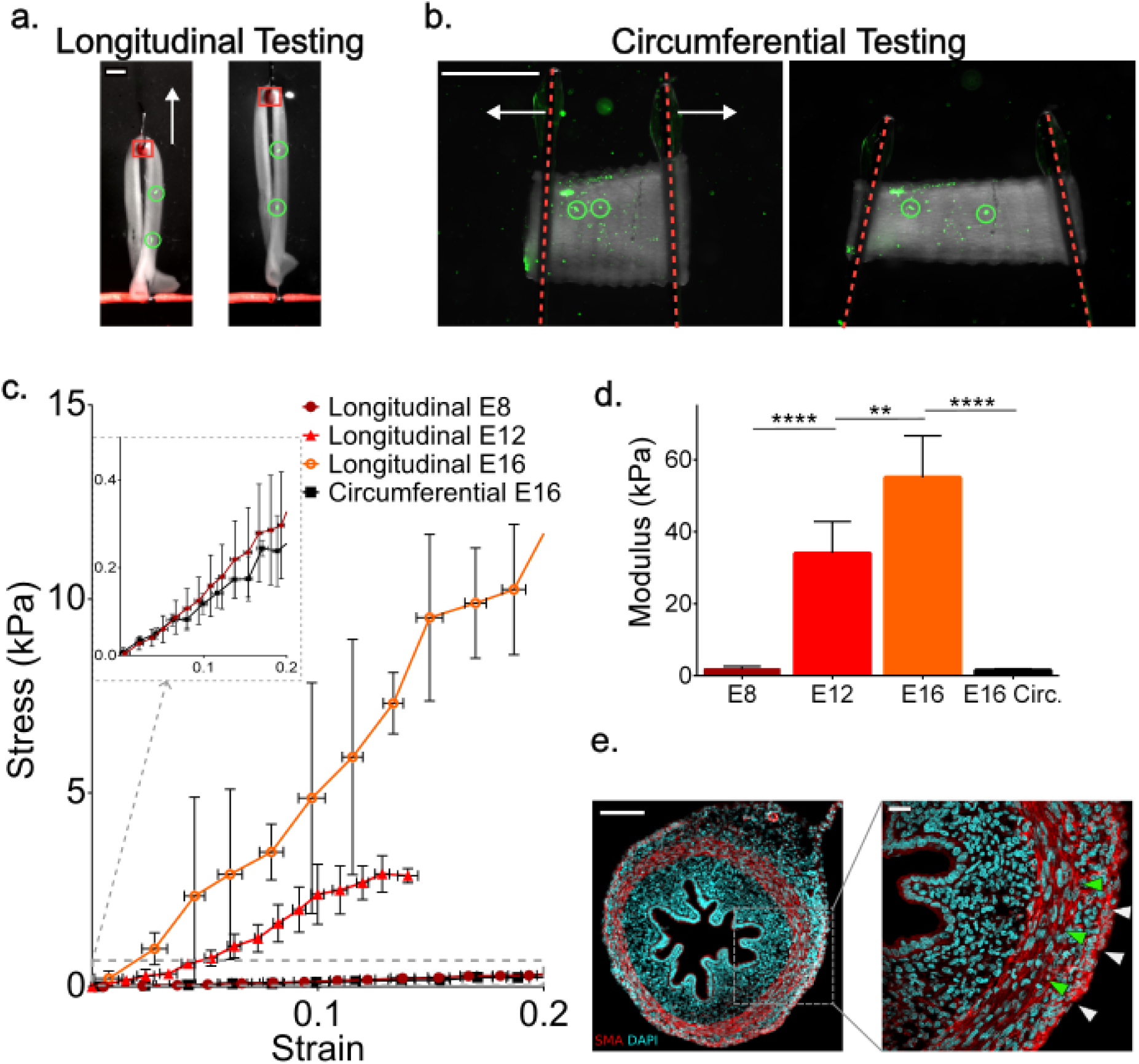
Tensile testing of the embryonic intestine reveals dynamic, anisotropic properties. (a) Images of a segment of E16 intestine before (left) and during (right) application of longitudinal stretch. Green circles indicate DiO tissue labels used to calculate strain. Red box marks the free end of the cantilever, which is hooked around the tube to apply stretch. White arrow indicates direction of cantilever displacement; scale bar = 1 mm. (b) A segment of intestine before (left) and during (right) circumferential tensile testing in a ring setup. Red dashed lines indicate tungsten rods bidirectionally displaced radially to stretch the circumference of the tube. Green circles indicate DiO tissue labels used to calculate strain. White arrow indicates direction of cantilever displacement; scale bar = 100 µm. (c) Mean longitudinal stress-strain curves for intestinal tissue at E8 (n=5), 12 (n=5), and 16 (n=6), and mean circumferential stress-strain curve at E16 (n=5); gray dashed box magnified in the inset to visualize longitudinal E8 and circumferential E16 groups. (d) Elastic modulus of samples represented in (b); *\*\*p <0.005; ****p <0.0001*. (e) Immunostaining for smooth muscle marker smooth muscle actin (SMA, red) and cell nuclei (DAPI, cyan) in E12 intestinal tube cross-section; gray box at left magnified at right with green and white arrowheads indicating circumferentially and longitudinally oriented smooth muscle layers, respectively; scale bar = 100 µm (left) and 50 µm (right).

### 2.3 Circumferential testing of intestinal tube mechanics

To quantify tensile properties in the circumferential direction, the above mechanical testing setup was modified to convert unidirectional displacement of the linear actuator into bi-directional movement of two tungsten beams in opposing directions as described previously [23]. As above, the device was situated on the stage of a Zeiss Axioxoom fluorescent macroscope to visualize tissue deformations and cantilever deflection. Samples were prepared by removal of the mesentery and excision of 1 mm long segments of intestine; a pair of tungsten cantilevers were inserted into the lumenal space, and fluorescent DiO markers affixed to the tissue as above. Displacement of the pair of cantilevers at a rate of 0.005 mm/s was used to stretch the ring of intestinal tissue, and strain was measured from a region equally spaced between the two cantilevers to minimize confounding boundary effects. Tracking was used as above to quantify strain and force, which was then used to construct stress-strain curves.

### 2.4 Fluorescence Microscopy

Chick intestinal tissue was fixed, cryosectioned, and immunostained as described previously [7]. Elastin(1:250, Millipore Sigma MAB2503), smooth muscle actin (1:250, Abcam ab5694), fibronectin (1:250, Developmental Studies Hybridoma Bank B3/D6), laminin (1:250, Abcam ab2034) were detected using fluorescently-conjugated secondary antibodies (1:500). Type I collagen (1:250; Abcam 34710) was visualized by amplification using a biotinylated secondary antibody (1:500, Jackson Immunoresearch 111-165-003) followed by incubation in streptavidin-HRP (Jackson Immunoresearch 016-030-084), and visualized with tyramide amplification (CF®555 Dye Tyramide; Biotium 96021). Samples were counterstained with DAPI, mounted in Fluoromount-G (00-4958-02; ThermoFisher Scientific), and coverslipped. Imaging was conducted on a Zeiss Axiozoom fluorescent macroscope or Zeiss LSM 880 laser scanning confocal microscope. For whole mount imaging of the elastic fiber network, fixed segments of intestinal tube were dehydrated in methanol, bleached, incubated with primary antibody for 3 days, secondary antibody for 2 days, and cleared with RapiClear (SunjinLab) [24].

### 2.5 Enzymatic Treatment

Enzymatic treatments to deplete elastin from the intestine were carried out following previously developed protocols [7]. Briefly, whole intestines (E8) or 2 cm long segments (E12 and 16) of intestinal tube and attached mesentery were rinsed in PBS, then incubated in either 0U/mL or 2U/mL of high purity porcine elastase (EC134, Elastin Products Company) for 4 hours at 37°C under light agitation. Off-target proteolysis and potential collagenolytic activity were limited by the inclusion of 0.1 mg/ml soybean trypsin inhibitor (Sigma-Aldrich) [25–27], confirmed by immunostaining for type I collagen (Supplemental Figure S1b).

### 2.6 Statistical Methods

All quantitative data were presented as mean +/± standard deviation. Statistical differences between groups were analyzed using one-way analysis of variance (ANOVA). A value of p < 0.05 represents the threshold for statistical significance; *ns : non-significant; *: (p>0.05); **: p <0.005; ****: p <0.0001*.

## 3. Results

### 3.1 Evolving axial mechanics of the developing intestinal tube

The stable equilibrium morphology of the buckled embryonic intestine depends on balancing the extensional energy of the mesentery with flexural stiffness of the intestinal tube [8]. Because prior efforts to understand the basis of mechanical properties during looping have focused primarily on the mesentery [14,22], here we sought to understand the determinants of material properties in the tube itself. To assess the evolving tensile properties of the intestinal tube along the longitudinal axis, we performed uniaxial tensile tests on a custom meso-scale mechanical tester as described previously [7,22]. In brief, a fine tungsten cantilever is used as a force transducer to load the tissue, and fluorescent markers are deposited onto the tissue to act as fiducial markers for strain analysis (Fig. 2a). The stress-strain behavior of the intestinal tube was approximately linear from E8 through E16 (Fig. 2c). However, the tensile modulus increased nearly 15-fold from the onset of gut looping at E8 to its completion at E16 (Fig. 2c, d). These dramatic changes in tissue stiffness were correlated with the emergence of an outer layer of smooth muscle cells encasing the tube and aligned along the longitudinal axis (Fig. 2e), raising the possibility that this outer longitudinal smooth muscle layer may contribute to the axial stiffness of the intestinal tube.

### 3.2 The embryonic small intestine is highly anisotropic

The mature intestine consists of three concentric layers of oriented smooth muscle: an inner longitudinal layer that lies within the submucosa, a thick band of circumferentially oriented smooth muscle that makes up much of the intestinal wall thickness, and a thin outer layer that also aligns longitudinally [28,29]. Based on the alignment of smooth muscle layers along alternating, orthogonal directions, we next asked whether this structural anisotropy translates to mechanical anisotropy as well. To test this, the mechanical tester was adapted to perform ring tests on segments of intestine, extracting tensile moduli associated with the circumferential direction (Fig. 2b). Owing to the small size of E8 and E12 intestines, these experiments were only carried out at E16, when looping ends. Despite the thick, aligned band of circumferential smooth muscle present, the circumferential modulus of E16 intestines was an order of magnitude lower than the longitudinal direction (Fig. 2d). This indicates that at the conclusion of looping, the embryonic intestine is characterized by dramatic mechanical anisotropy.

### 3.3 Longitudinal stiffening of the Intestine is correlated with ECM deposition

We were surprised to observe that the intestinal modulus is higher in the longitudinal direction, where only a thin layer of smooth muscle is aligned with the loading direction, when compared to the circumferential direction, which coincides with a thick layer of mature smooth muscle (Fig. 2e). This led us to consider that the ECM, rather than smooth muscle cells, may dictate the mechanical properties of the intestine. Immunostaining for common ECM components between E8, when looping initiates, and E16, when looping is complete, revealed dynamic composition and organization of matrix components as looping proceeds (Fig. 3). Fibronectin increased in the tube over time, and appeared highly aligned parallel to each smooth muscle layer (Fig. 3a). As expected, laminin was strongly expressed in the basement membrane associated with the mucosal epithelium (Fig. 3b). However, abundance in the basement membrane diminished over time. Interestingly, laminin, which primarily functions as a basement membrane protein, was strongly associated with the smooth muscle layers, sharing their alignment in a similar manner to fibronectin. Type I collagen also increased in abundance and became closely associated with smooth muscle layers as development proceeds (Fig. 3c). Finally, based on the importance of elastic fibers for establishing the mechanical properties of its attached mesentery [7], we examined elastin organization within the intestinal tube as well (Fig. 4). Elastin staining differed from other ECM components in that it was abundantly expressed in the outer longitudinal layer of the tube, but only sparsely present in the large circumferential smooth muscle layer (Fig. 4a). 3-D imaging of optically cleared intestines revealed an intricate network of radially oriented elastic fibers within the circumferential smooth muscle layer, wrapping around individual muscle bundles (Fig. 4b), but little to no elastin oriented circumferentially within the smooth muscle layer. Therefore, elastin is enriched in the longitudinal smooth muscle layer, which aligns with the direction of high tensile modulus, but diminished in the circumferential layer, which is associated with a much lower modulus. The correlation of elastin orientation and abundance with modulus led us to next consider whether elastin - rather than oriented smooth muscle cells - plays a central role in establishing mechanical anisotropy of the intestinal tube.

**Figure 3:**
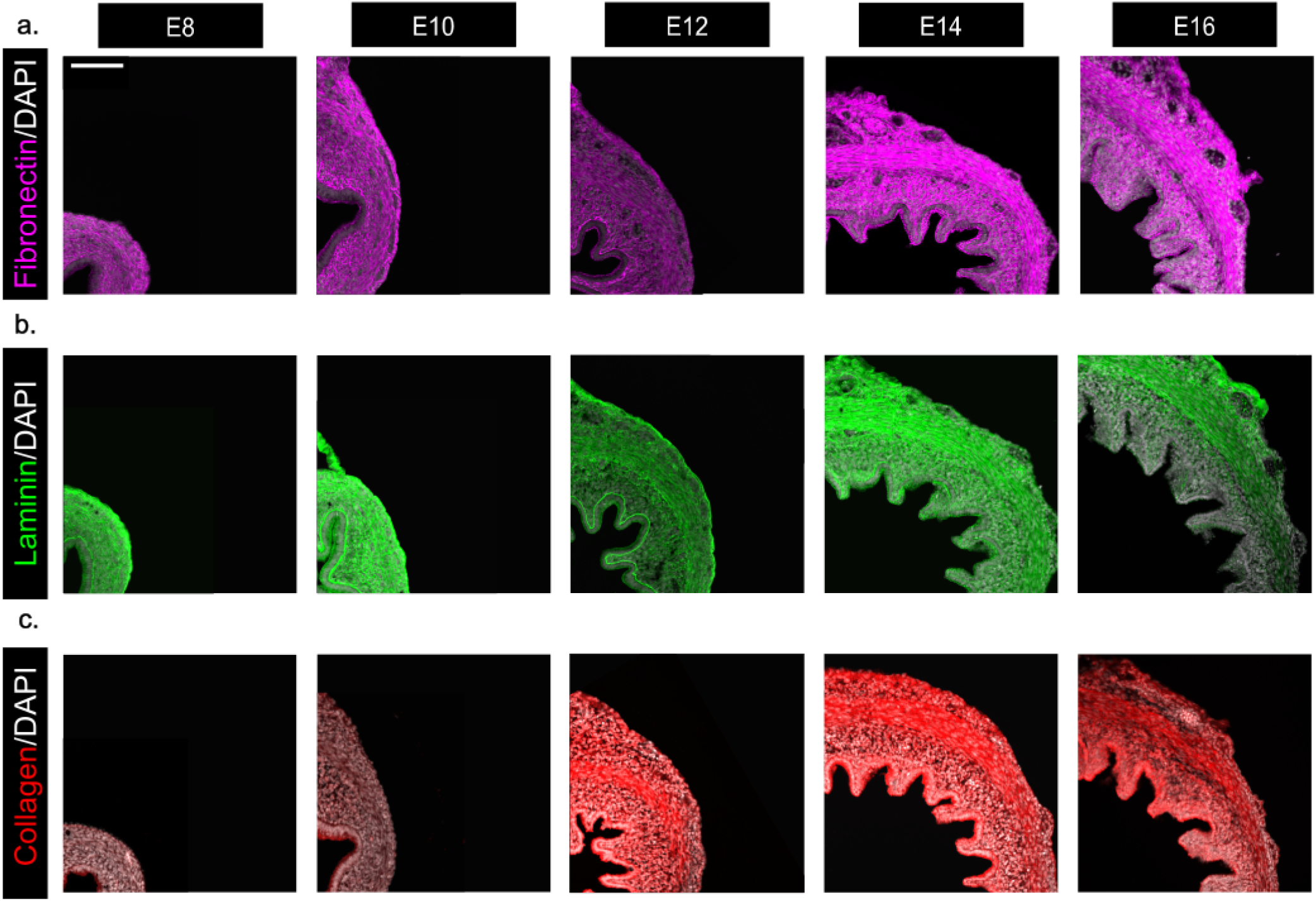
Time course of ECM deposition in the intestinal tube. Immunostaining of ECM proteins fibronectin (a, magenta), laminin (b, green), and collagen (c, red) at embryonic stages E8, 10, 12, 14, and 16, counterstained with DAPI to visualize cell nuclei (gray); embryonic dorsal direction is upward. Scale bar = 100 µm.

**Figure 4:**
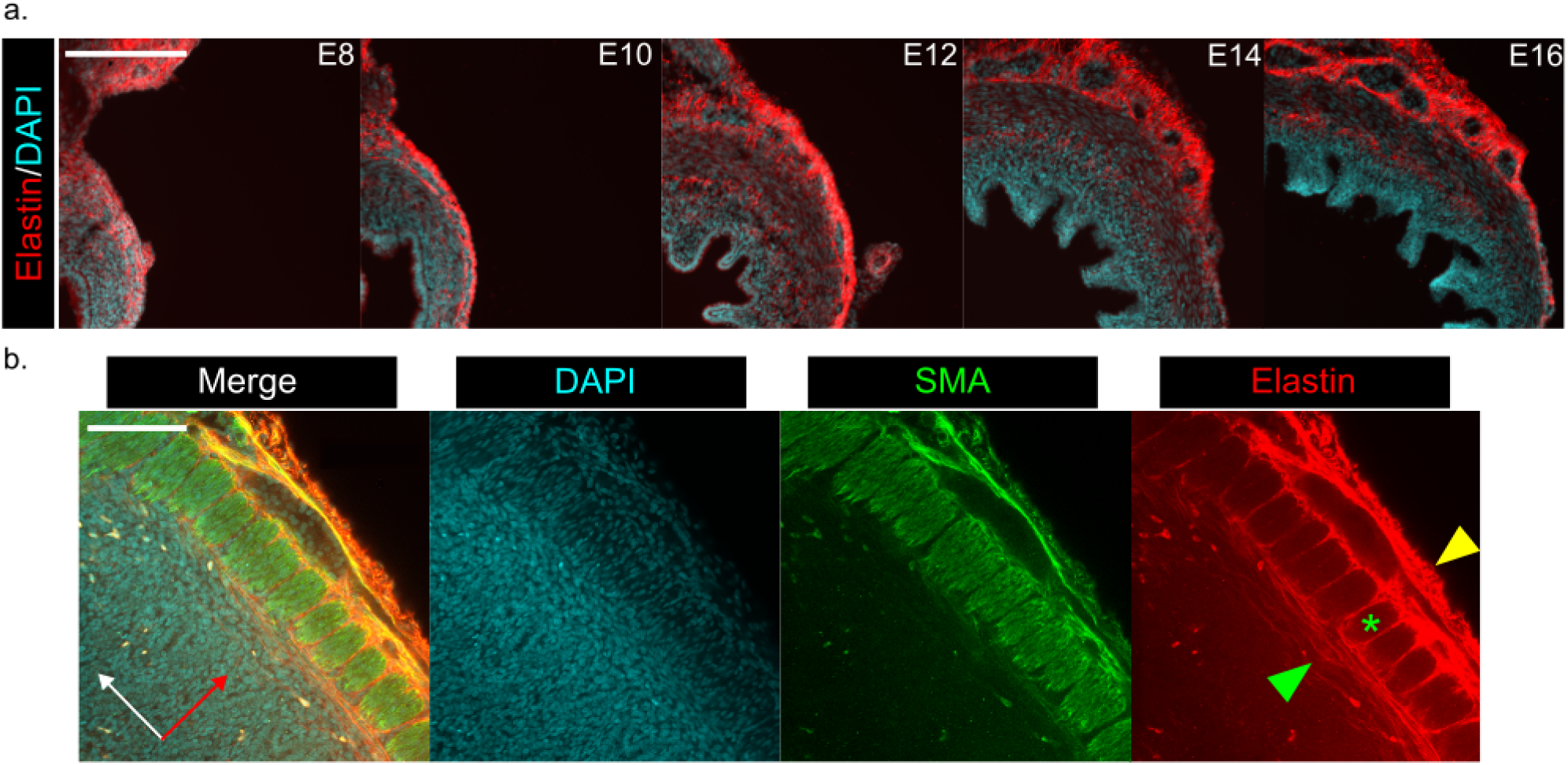
Elastin organization in the gut tube is shown to align with smooth muscle layers. (a) Time course of elastin in the gut tube. Dorsal quadrant of the gut tube is pictured with dorsal oriented up. Scale bar = 200 µm. (b) Sagittal cross-section of an E16 gut tube stained for elastin, where white and red indicate the longitudinal and radial axes of the intestine, respectively. Immunostaining for smooth muscle actin (SMA, green) and counterstaining for nuclei (DAPI, cyan) reveals elastin wrapping around the circumferential smooth muscle bundles (green asterisk) as well as aligning with the inner (green arrowhead) and outer (yellow arrowhead) longitudinal muscle layers. Scale bar = 100 µm.

### 3.4 Elastic fibers are responsible for anisotropy of the embryonic small intestine

To test whether elastic fibers in the intestinal tube contribute to tissue anisotropy and mechanics, we performed selective enzymatic disruption of elastin using elastase [7]. Effective degradation of elastic fibers in the intestinal tube was confirmed by immunostaining (Supplementary Fig. 1a). Following elastase treatment, we performed tensile tests to examine how elastin depletion influences longitudinal stiffness of the intestine across developmental stages. At E8, when elastin in the tube is disorganized and diffusely distributed through the mesenchyme and the outer longitudinal smooth muscle layer has not yet formed, elastase treatment had no effect on the modulus (Fig. 5a, d). However, at E12 and E16, when elastic fibers are more defined and tightly packaged in the outer longitudinal muscle layer, we observed dramatic reductions in modulus with elastase treatment, by as much as 95% (Fig. 5b-d). These results suggest that elastin is a key matrix component in defining the longitudinal properties of the intestinal tube, which in turn dictates bending stiffness of the tube during looping morphogenesis. We next examined whether elastase treatment similarly alters circumferential properties, focusing on E16, when such tests are possible. Unexpectedly, circumferential properties were unaltered by elastase treatment (Fig. 6). As a result, elastase treatment largely removed mechanical anisotropy in the tube: longitudinal and circumferential moduli were nearly identical after elastase treatment, despite their separation by an order of magnitude in controls. These data suggest that elastic fibers confer anisotropy and longitudinal stiffness to the developing intestine.

**Figure 5:**
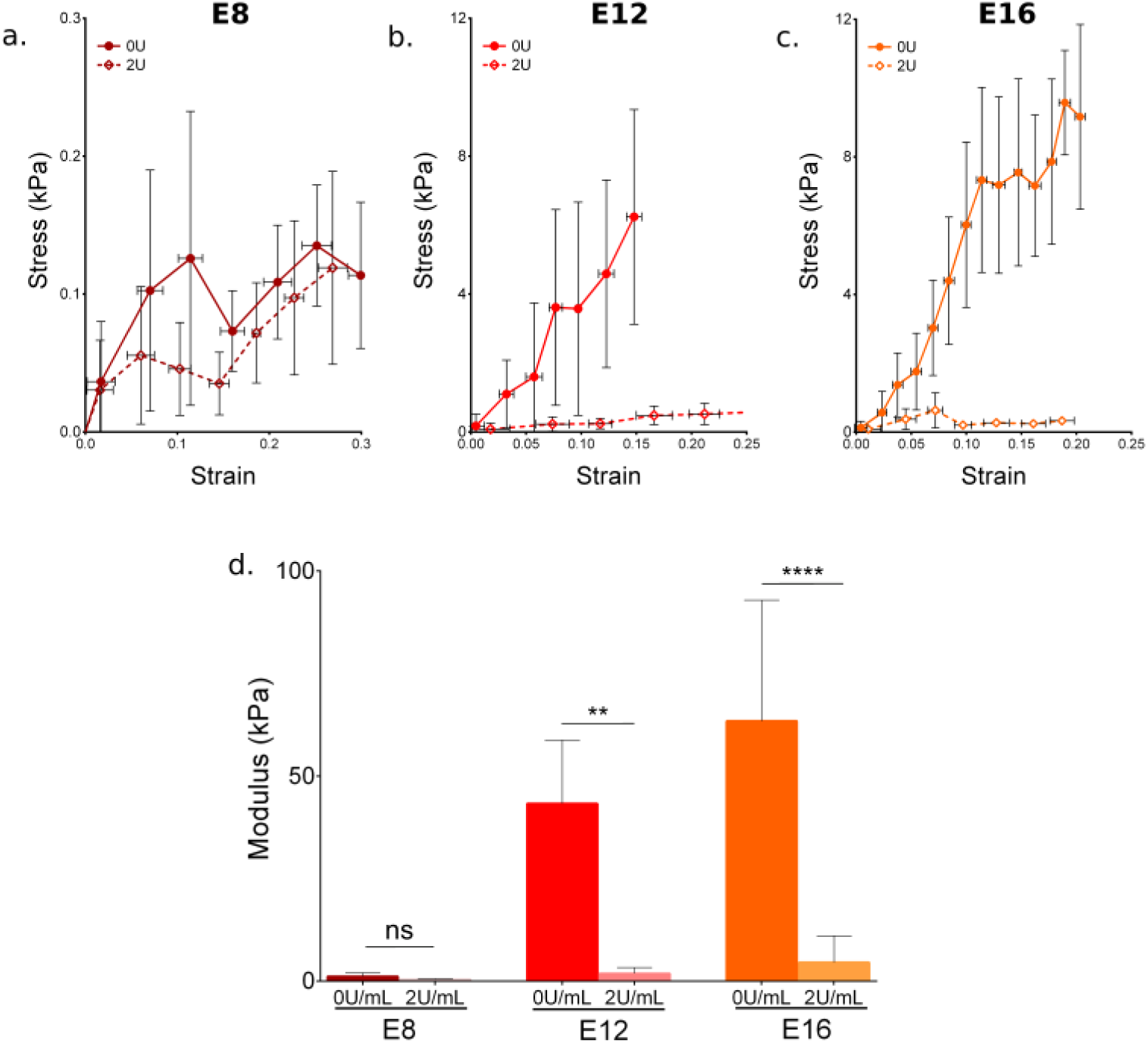
Effects of elastase treatment on longitudinal properties of the intestinal tube. (a-c) Mean longitudinal stress-strain curves of E8 (a), E12 (b), and E16 (c) intestines following treatment with for 0 U/mL (solid symbols/lines) and 2 U/mL elastase (open symbols/dashed lines); n=6 for E16 0 U/mL, and n=5 for all other groups. (d) Elastic modulus of intestines represented in (a-c); *ns: non-significant; **: p<0.005, ****: p<0.0001*.

**Figure 6:**
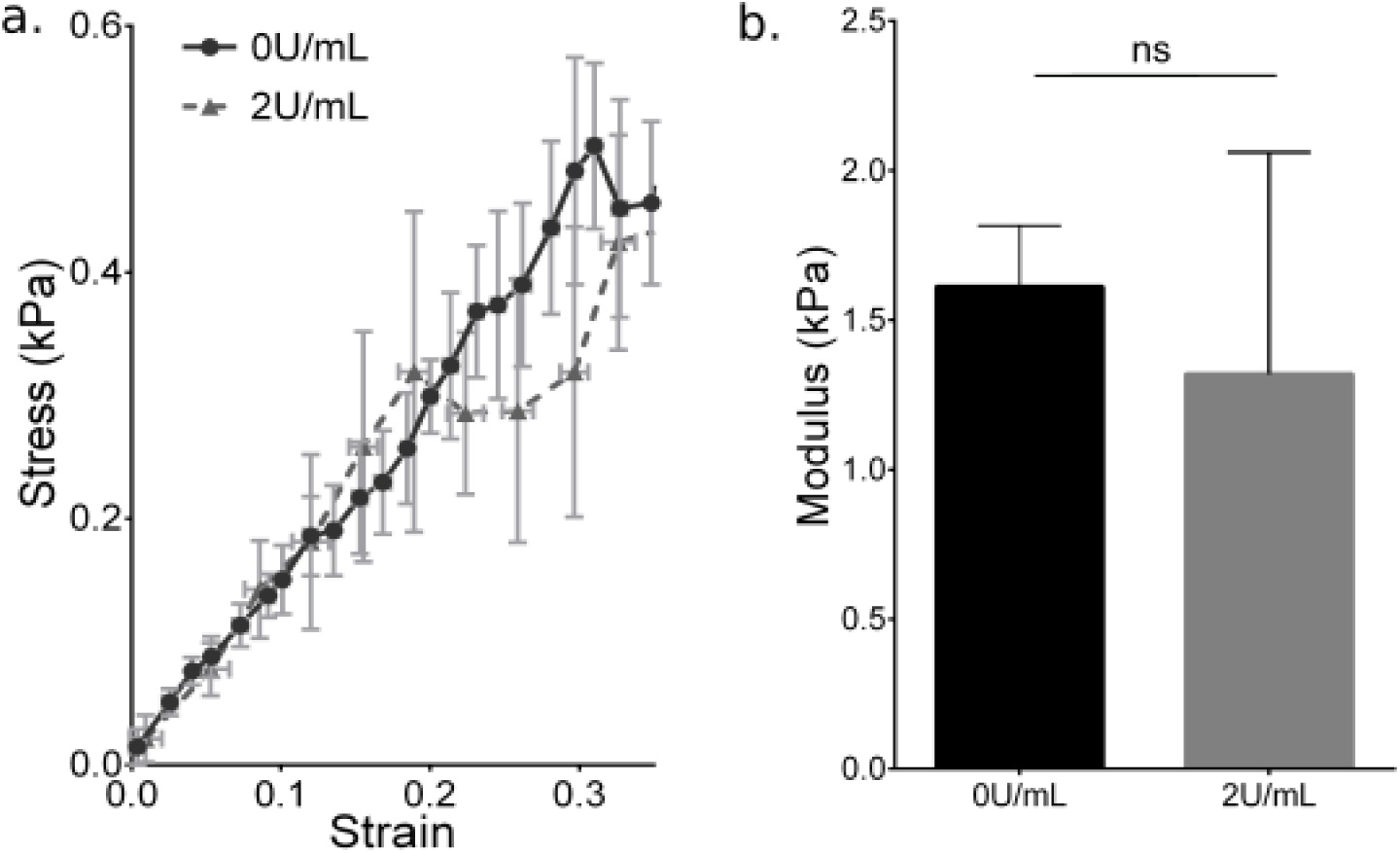
Effects of elastase treatment on circumferential properties of the intestinal tube. (a) Mean circumferential stress-strain curves of E16 intestinal tube following treatment with 0U/mL (black circle, solid line, n = 5) and 2U/mL elastase (gray triangle, dashed line, n =5). (b) Elastic modulus of E16 intestines represented in (a); *ns: non-significant*.

## 4. Discussion

In the present study, we investigated material properties of the embryonic small intestine during looping morphogenesis, when these properties must be precisely coordinated with growth and stiffness of the attached mesentery to achieve proper packing and placement of the intestine within the body cavity. We observed that the longitudinal stiffness of the intestinal tube increases significantly from the onset of looping at E8 to its completion at E16 (Fig. 2). At E16, the tube is highly anisotropic, with an order of magnitude separating the modulus in the stiffer longitudinal direction from that of the softer circumferential direction. Combining histological analyses with enzymatic treatments and mechanical testing, we reveal that this anisotropy is due to elastic fibers enriched in the outer longitudinal smooth muscle layer, establishing bending stiffness of the intestinal tube during looping. These findings enhance our understanding of a critical step in gastrointestinal development, and may contribute to a broader reframing of congenital disorders in part as errors in the embryonic establishment of tissue material properties.

The biomechanical role of elastin has mostly been studied in adult tissues, with particular focus on cardiovascular and orthopaedic tissues [27,30–33]. However, we recently observed an important role for elastin during gut looping, where mesentery stiffness was largely dependent on the content of elastic fibers in the tissue [7]. This was somewhat unexpected, as elastin is most frequently considered for its role in tissues that undergo large cyclic deformations such as arteries and ligaments, leading to the idea that elastin may have distinct roles during embryogenesis and postnatally. In the present study, we again find an important role for elastin in morphogenesis, but here, elastin in the intestinal tube was closely associated with smooth muscle differentiation, suggesting that it may have important postnatal function independent of its role in looping morphogenesis. For example, radially oriented fibers that interdigitate with bundles of circumferential smooth muscle (Fig. 4b) may be passively stretched during peristalsis, and function to enable recoil to a dilated lumen morphology following contraction. Similarly, close association of elastin with the outer longitudinal smooth muscle layer points to an important and previously unappreciated potential role for elastin in gut motility: rhythmic peristaltic contractions that drive contents through the intestine result in large deformations that must be cyclically reversed when the muscle relaxes, and it is possible that elastic fibers perform this function. This highlights an intriguing puzzle in the evolution of developmental mechanisms: during embryogenesis, the goals of shaping a tissue and establishing its postnatal function must be achieved concomitantly, despite having differing needs from a material property standpoint. For example, although elastic fibers in the tube likely function in stabilizing and reversing peristaltic contractions, they also are responsible for over 90% of the bending stiffness of the intestinal tube. This represents an energetic barrier to looping, as the mesentery must be sufficiently stiff and thick, and the growth differential between these tissues sufficiently large as to generate compressive forces large enough to drive buckling. Tissue geometry, growth rate, and stiffness are each complex multi-genic traits, and the coordination between these traits in the mesentery with elastic fiber deposition in the tube - which is likely dictated by physiologic needs of the tissue postnatally - is consistent with the notion of mechanical feedback, wherein cell behaviors of the mesentery are under some level of control from tension in the tissue, which depends on tube stiffness.

Mechanics has been demonstrated to have important roles during intestinal development, both as an effector of developmental programs to shape tissues [9,28,29,34] and as an upstream regulator of cell behaviors [35], such as placement and orientation of smooth muscle layers within the intestine [12,36]. Concurrent with looping, a second buckling event is initiated on the inner luminal surface of the avian intestine, wherein the initially smooth endoderm-derived epithelium that lines the lumen is first buckled into longitudinal ridges, which then through a second buckling event form zigzags that presage the outgrowth of villi from their inflection points. In this context, buckling is driven by rapid growth of the inner endodermal epithelium against the constraint of oriented mesodermal smooth muscle layers - first the circumferential layer which drives buckling into longitudinal ridges, then the longitudinal smooth muscle layer, which buckles those ridges into zigzags [28]. The function of smooth muscle as stiff constraints shaping buckling epithelia is also observed in lung branching [17]. However, our results suggest that in the intestine, the mechanical constraint driving buckling of ridges into zigzags may not be the longitudinal smooth muscle itself, but the elastin-rich extracellular matrix that localizes within this layer and accounts for over 90% of the tube’s stiffness in the longitudinal direction. To test this, we examined the effects of elastase on epithelial morphology inside the tube, but found that elastase treated intestines retain their newly formed zigzag morphology (data not shown). This suggests that, if elastin is involved in villi morphogenesis, it may not be needed to maintain tissue organization once the process is complete.

As the importance of elastin for morphogenesis and function of the small intestine becomes increasingly apparent, it will be valuable to understand the upstream biochemical and physical cues that regulate elastic fiber deposition and patterning as well. However, relatively little is known on the upstream factors controlling patterning of elastic fibers in the intestine. TGF-β signaling has recently been implicated in deposition of other ECM components in the developing intestine [9], and mutations in Latent TGF-β Binding Proteins (LTBP) are associated with defective elastic fiber assembly in the intestine [37], suggesting that this may be an important pathway in patterning elastic fiber formation. Owing to the spatio-temporal coordination between smooth muscle organization and elastin-rich ECM, it is possible that both are patterned through the same upstream regulatory cues. Radial placement of smooth muscle layers within the intestine results from an activator-inhibitor type interaction between Hedgehog and BMP ligands [36]. In future work, it will be informative to examine whether the same pathways regulate elastic fiber patterning by experimentally shifting spatial gradients of Hedgehog and BMP and examining the effects on elastin expression. While it is possible that elastic fiber assembly is downstream of smooth muscle differentiation, the distinct patterns of elastic fiber deposition observed in circumferential and longitudinal muscle layers suggest the relationship may be more complicated.

Finally, several human syndromes have been traced to mutations *ELN*, the gene encoding elastin [38]. While the most life threatening complications tend to be cardiovascular in nature, several include defects of the gastrointestinal tract as well. Cutis laxa, a syndrome resulting from mutations of the elastin gene ELN, results in multiple intestinal diverticula [39]. Williams-Beuren Syndrome results from a microdeletion including part of the coding domain of *ELN* along with several other genes, and results in several complications of the gastrointestinal tract, including diverticulitis, severe constipation, hematochezia, and diarrhea [40]. These and other human genetic disorders of the elastic fiber network are consistent with an important mechanical role for elastic fibers in function of the small intestine, and highlight the clinical importance of understanding the developmental basis of this key mechanical component of the intestine.

## Acknowledgements

We thank members of the Nerurkar Lab for their valuable scientific input. This work was funded by NIDDK grant R01DK131236 (N.L.N.) with additional support from the Columbia University Digestive and Liver Disease Research Center (1P30DK132710).

## References

[1] Y.C. Fung, Biomechanics: Mechanical Properties of Living Tissues, Second Edition, Springer, 1993.

[2] J. Crest, A. Diz-Muñoz, D.Y. Chen, D.A. Fletcher, D. Bilder, Organ sculpting by patterned extracellular matrix stiffness, eLife 6 (2017). 10.7554/eLife.24958.

[3] A. Sivakumar, A. Mahadevan, M.E. Lauer, R.J. Narvaez, S. Ramesh, C.M. Demler, N.R. Souchet, V.C. Hascall, R.J. Midura, S. Garantziotis, D.B. Frank, K. Kimata, N.A. Kurpios, Midgut Laterality Is Driven by Hyaluronan on the Right, Dev. Cell 46 (2018) 533–551.e5. 10.1016/j.devcel.2018.08.002.

[4] C. Kyprianou, N. Christodoulou, R.S. Hamilton, W. Nahaboo, D.S. Boomgaard, G. Amadei, I. Migeotte, M. Zernicka-Goetz, Basement membrane remodelling regulates mouse embryogenesis, Nature 582 (2020) 253–258. 10.1038/s41586-020-2264-2.

[5] A. Mongera, P. Rowghanian, H.J. Gustafson, E. Shelton, D.A. Kealhofer, E.K. Carn, F. Serwane, A.A. Lucio, J. Giammona, O. Campàs, A fluid-to-solid jamming transition underlies vertebrate body axis elongation, Nature 561 (2018) 401–405. 10.1038/s41586-018-0479-2.

[6] J. Zhou, H.Y. Kim, L.A. Davidson, Actomyosin stiffens the vertebrate embryo during crucial stages of elongation and neural tube closure., Dev. Camb. Engl. 136 (2009) 677–88. 10.1242/dev.026211.

[7] E.A. Loffet, J.F. Durel, J. Gao, R. Kam, H. Lim, N.L. Nerurkar, Elastic fibers define embryonic tissue stiffness to enable buckling morphogenesis of the small intestine, Biomaterials 303 (2023) 122405. 10.1016/j.biomaterials.2023.122405.

[8] T. Savin, N. a Kurpios, A.E. Shyer, P. Florescu, H. Liang, L. Mahadevan, E. Al., On the growth and form of the gut, Nature 476 (2011) 57–62.

[9] H.K. Gill, S. Yin, N.L. Nerurkar, J.C. Lawlor, T.R. Huycke, L. Mahadevan, C.J. Tabin, Hox genes modulate physical forces to differentially shape small and large intestinal epithelia, (2023) 2023.03.15.532602. 10.1101/2023.03.15.532602.

[10] T. Tallinen, J.Y. Chung, F. Rousseau, N. Girard, J. Lefèvre, L. Mahadevan, On the growth and form of cortical convolutions, Nat. Phys. 12 (2016) 588–593.

[11] C.M. Nelson, On Buckling Morphogenesis, J. Biomech. Eng. 138 (2016) 021005. 10.1115/1.4032128.

[12] J.F. Durel, N.L. Nerurkar, Mechanobiology of vertebrate gut morphogenesis, Curr. Opin. Genet. Dev. 63 (2020) 45–52. 10.1016/j.gde.2020.04.002.

[13] F.I. Luks, Anomalies of intestinal rotation, in: Fundam. Pediatr. Surg., Springer New York, New York, NY, 2011: pp. 373–380. 10.1007/978-1-4419-6643-8_48.

[14] E.A. Loffet, J.F. Durel, N.L. Nerurkar, Evo-Devo Mechanobiology: The Missing Link, Integr. Comp. Biol. (2023) icad033. 10.1093/icb/icad033.

[15] N.L. Nerurkar, C.H. Lee, L. Mahadevan, C.J. Tabin, Molecular control of macroscopic forces drives formation of the vertebrate hindgut, Nature 565 (2019) 480–484. 10.1038/s41586-018-0865-9.

[16] J.W. Spurlin, M.J. Siedlik, B.A. Nerger, M.F. Pang, S. Jayaraman, R. Zhang, C.M. Nelson, Mesenchymal proteases and tissue fluidity remodel the extracellular matrix during airway epithelial branching in the embryonic avian lung, Dev. Camb. 146 (2019). 10.1242/dev.175257.

[17] K. Goodwin, B. Lemma, P. Zhang, A. Boukind, C.M. Nelson, Plasticity in airway smooth muscle differentiation during mouse lung development, Dev. Cell 58 (2023) 338–347.e4. 10.1016/j.devcel.2023.02.002.

[18] S. Yang, K.H. Palmquist, L. Nathan, C.R. Pfeifer, P.J. Schultheiss, A. Sharma, L.C. Kam, P.W. Miller, A.E. Shyer, A.R. Rodrigues, Morphogens enable interacting supracellular phases that generate organ architecture, Science 382 (2023) eadg5579. 10.1126/science.adg5579.

[19] E.H. Barriga, K. Franze, G. Charras, R. Mayor, Tissue stiffening coordinates morphogenesis by triggering collective cell migration in vivo, Nature 554 (2018) 523–527. 10.1038/nature25742.

[20] L. Legoff, H. Rouault, T. Lecuit, A global pattern of mechanical stress polarizes cell divisions and cell shape in the growing Drosophila wing disc., Dev. Camb. Engl. 140 (2013) 4051–9. 10.1242/dev.090878.

[21] N. Desprat, W. Supatto, P.A. Pouille, E. Beaurepaire, E. Farge, Tissue Deformation Modulates Twist Expression to Determine Anterior Midgut Differentiation in Drosophila Embryos, Dev. Cell 15 (2008) 470–477. 10.1016/j.devcel.2008.07.009.

[22] N.L. Nerurkar, L. Mahadevan, C.J. Tabin, BMP signaling controls buckling forces to modulate looping morphogenesis of the gut, Proc. Natl. Acad. Sci. U. S. A. 114 (2017) 2277–2282. 10.1073/pnas.1700307114.

[23] P. Oikonomou, L. Calvary, H.C. Cirne, A.E. Welch, J.F. Durel, O. Powell, N.L. Nerurkar, Application of tissue-scale tension to avian epithelia in vivo to study multiscale mechanical properties and inter-germ layer coupling, BioRxiv Prepr. Serv. Biol. (2024) 2024.04.04.588089. 10.1101/2024.04.04.588089.

[24] G. Timin, M.C. Milinkovitch, High-resolution confocal and light-sheet imaging of collagen 3D network architecture in very large samples, iScience 26 (2023) 106452. 10.1016/j.isci.2023.106452.

[25] Y.F. Missirlis, Use of enzymolysis techniques in studying the mechanical properties of connective tissue components, J. Bioeng. 1 (1977) 215–222.

[26] H. Oxlund, J. Manschot, A. Viidik, The role of elastin in the mechanical properties of skin, J. Biomech. 21 (1988) 213–218. 10.1016/0021-9290(88)90172-8.

[27] H.B. Henninger, C.J. Underwood, S.J. Romney, G.L. Davis, J.A. Weiss, Effect of elastin digestion on the quasi-static tensile response of medial collateral ligament, J. Orthop. Res. 31 (2013) 1226–1233. 10.1002/jor.22352.

[28] A.E. Shyer, T. Tallinen, N.L. Nerurkar, Z. Wei, E.S. Gil, D.L. Kaplan, C.J. Tabin, L. Mahadevan, Villification: how the gut gets its villi, Science 342 (2013) 212–218. 10.1126/science.1238842.

[29] N.R. Chevalier, Physical organogenesis of the gut, Development 149 (2022) dev200765. 10.1242/dev.200765.

[30] H. Trębacz, A. Barzycka, Mechanical Properties and Functions of Elastin: An Overview, Biomolecules 13 (2023) 574. 10.3390/biom13030574.

[31] D.Y. Li, B. Brooke, E.C. Davis, R.P. Mecham, L.K. Sorensen, B.B. Boak, E. Eichwald, M.T. Keating, Elastin is an essential determinant of arterial morphogenesis., Nature 393 (1998) 276–280. 10.1038/30522.

[32] J.E. Wagenseil, R.P. Mecham, Elastin in large artery stiffness and hypertension, J Cardiovasc. Transl. Res. 5 (2012) 264–273. 10.1007/s12265-012-9349-8.

[33] J.E. Wagenseil, N.L. Nerurkar, R.H. Knutsen, R.J. Okamoto, D.Y. Li, R.P. Mecham, Effects of elastin haploinsufficiency on the mechanical behavior of mouse arteries., Am. J. Physiol. Heart Circ. Physiol. 289 (2005) H1209–17. 10.1152/ajpheart.00046.2005.

[34] H.K. Gill, S. Yin, J.C. Lawlor, T.R. Huycke, N.L. Nerurkar, C.J. Tabin, L. Mahadevan, The developmental mechanics of divergent buckling patterns in the chick gut, Proc. Natl. Acad. Sci. 121 (2024) e2310992121. 10.1073/pnas.2310992121.

[35] C. Collinet, T. Lecuit, Programmed and self-organized flow of information during morphogenesis, Nat. Rev. Mol. Cell Biol. 22 (2021) 245–265. 10.1038/s41580-020-00318-6.

[36] T.R. Huycke, B.M. Miller, H.K. Gill, N.L. Nerurkar, D. Sprinzak, M. L., C.J. Tabin, Genetic and mechanical regulation of intestinal smooth muscle development, Cell (n.d.).

[37] Z. Urban, V. Hucthagowder, N. Schürmann, V. Todorovic, L. Zilberberg, J. Choi, C. Sens, C.W. Brown, R.D. Clark, K.E. Holland, M. Marble, L.Y. Sakai, B. Dabovic, D.B. Rifkin, E.C. Davis, Mutations in LTBP4 cause a syndrome of impaired pulmonary, gastrointestinal, genitourinary, musculoskeletal, and dermal development, Am. J. Hum. Genet. 85 (2009) 593–605. 10.1016/j.ajhg.2009.09.013.

[38] M.L.D. Lasio, B.A. Kozel, Elastin-Driven Genetic Diseases, Matrix Biol. J. Int. Soc. Matrix Biol. 71–72 (2018) 144–160. 10.1016/j.matbio.2018.02.021.

[39] A. Damkier, F. Brandrup, H. Starklint, Cutis laxa: autosomal dominant inheritance in five generations, Clin. Genet. 39 (1991) 321–329. 10.1111/j.1399-0004.1991.tb03038.x.

[40] M. Boechler, Y.-P. Fu, N. Raja, E. Ruiz-Escobar, L. Nimmagadda, S. Osgood, M.D. Levin, C. Hadigan, B.A. Kozel, Gastrointestinal manifestations in Williams syndrome: A prospective analysis of an adult and pediatric cohort, Am. J. Med. Genet. A. n/a (n.d.) e63827. 10.1002/ajmg.a.63827.

