## Supplemental Figure 1 for "Material Properties Of The Embryonic Small Intestine During Buckling Morphogenesis"

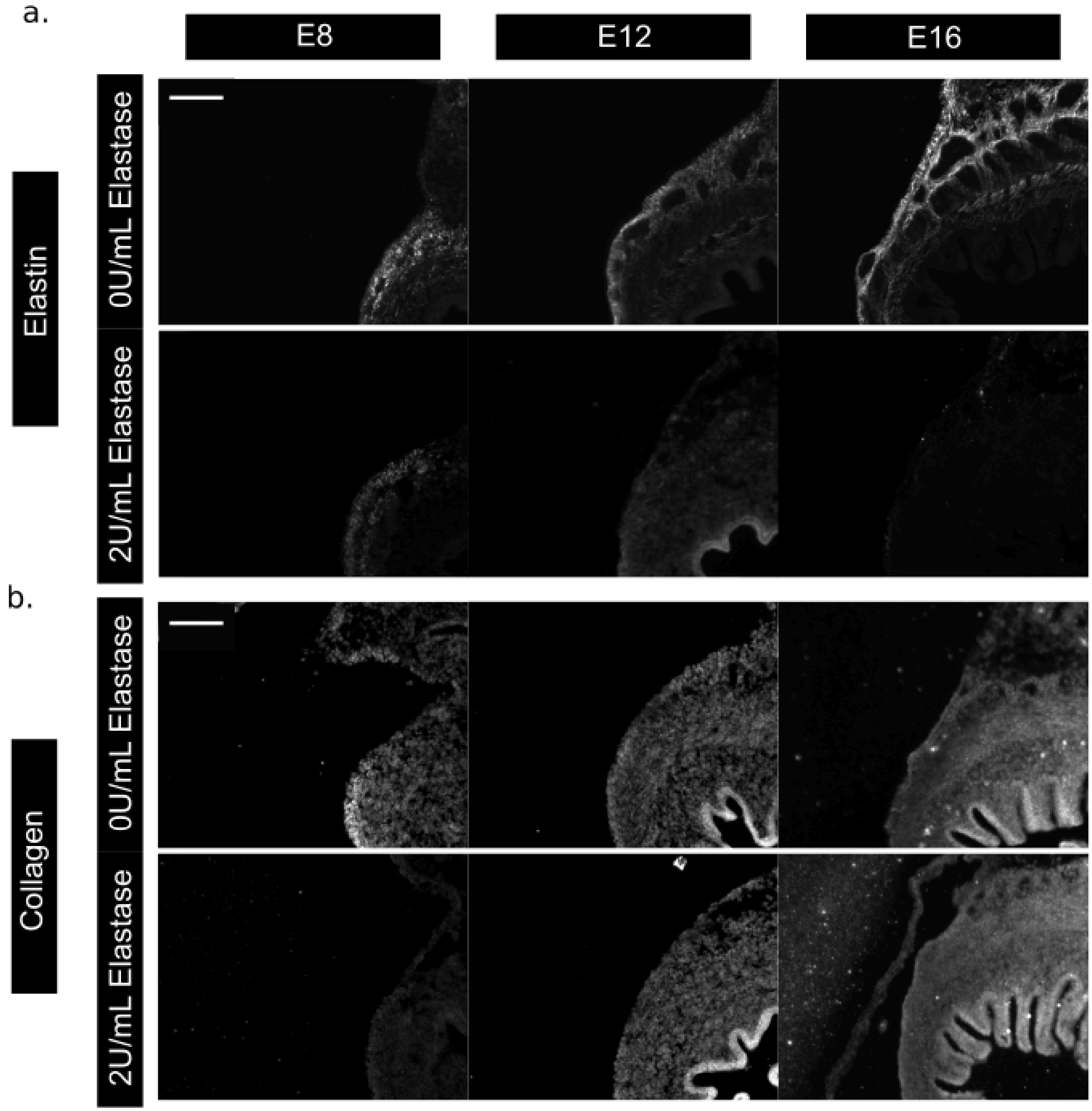


**Supplemental Figure 1: Elastase treatment depletes elastin from the intestinal tube with minimal effects on collagen** (a) Immunostaining of elastin (gray) in cross-sections of E8, E12, and E16 intestines following treatment with 0 U/mL (top) or 2 U/mL elastase. Scale bar = 100 μm. (b) Immunostaining of collagen (gray) in cross-sections of E8, E12, and E16 intestines following treatment with 0 U/mL (top) or 2 U/mL elastase. Scale bar = 100 μm.
